# Origins and Evolution of Novel *Bacteroides* in Captive Apes

**DOI:** 10.1101/2023.10.20.563286

**Authors:** Alexandra H. Nishida, Howard Ochman

## Abstract

Bacterial strains evolve in response to the gut environment of their hosts, with genomic changes that influence their interactions with hosts as well as with other members of the gut community. Great apes in captivity have acquired strains of *Bacteroides xylanisolvens*, which are common within gut microbiome of humans but not typically found other apes, thereby enabling characterization of strain evolution following colonization. Here, we isolate, sequence and reconstruct the history of gene gain and loss events in numerous captive-ape-associated strains since their divergence from their closest human-associated strains. We show that multiple captive-ape-associated *B. xylanisolvens* lineages have independently acquired gene complexes that encode functions related to host mucin metabolism. Our results support the finding of high genome fluidity in *Bacteroides*, in that several strains, in moving from humans to captive apes, have rapidly gained large genomic regions that augment metabolic properties not previously present in their relatives.

**Significance statement:** Chronicling the changes that occur in bacterial genomes after a host-switch event is normally difficult due to age of most bacteria-host associations, which renders uncertainties about the bacterial ancestor (and ancestral genome) prior to colonization of the new host. However, the gut microbiomes of great apes in captivity contain bacterial strains that are unique to humans, allowing fine-scale assessment and reconstruction of the genomic changes that follow colonization. By sequencing and comparing closely related strains of *Bacteroides* that are restricted both to human and to captive great apes, we found that multiple bacterial lineages convergently acquired sets of genes involved in the metabolism of dietary polysaccharides. These results show that over relatively short timescales, the incorporation of strains into microbiomes involves large-scale genomic events that correspond to characteristics of the new host environment.

## Introduction

Characterization of the compositional features of the gut microbial communities inhabiting humans and other great apes has shown that several bacterial taxa, such as *Bacteroides* and *Faecalibacterium*, are characteristic of humans but rarely observed in wild apes (Moeller et al. 2014; Hicks et al. 2018; Campbell et al. 2020). In captivity, however, the gut microbiome compositions of great ape species converge through the loss of bacterial strains unique to each wild-ape species and the acquisition of bacterial strains otherwise restricted to humans (Houtz et al. 2021; Nishida and Ochman 2021). In particular, the human-associated strains of *Bacteroides* that now reside in captive apes can divulge how bacterial genomes have evolved following host-switch events.

*Bacteroides* species have the ability to catabolize complex plant polysaccharides, which are indigestible by humans, into short-chain fatty acids that are consumable by hosts and other microbes (Briggs et al. 2021; Pruss et al 2021). Additionally, the ability of some *Bacteroides* strains to breakdown host-derived colonic mucosal glycans and human milk oligosaccharides enables them to outcompete other bacterial species and engraft in a community (Marcobal et al. 2011; Feng et al 2021). Recent work has demonstrated that individual *Bacteroides* species tend to specialize in degrading either host-derived or dietary-derived polysaccharides, suggesting that species can co-exist because they occupy distinct functional niches [10]. In contrast to other *Bacteroides* species, *Bacteroides xylanisolvens* strains exhibit extreme variation in the ability to metabolize host mucin and plant-derived polysaccharides (Lewis et al. 2020; Hao et al. 2021; Pudlo et al. 2021). Moreover, these functional differences are evident among strains that are otherwise identical by ribosomal RNA profiling (Pudlo et al. 2021).

Gene content differences among *Bacteroides* emerge through a combination of mechanisms including mutation (Groussin et al. 2021), duplication (Schofield et al. 2018), homologous recombination (Pudlo et al. 2021), phage transduction (Ogilivie et al. 2012; Campbell et al. 2020) and integrative conjugative elements (Hussain et al. 2017). Previous studies have followed strain evolution of *Bacteroides* species within hosts on a time scale of weeks to months by examining single-nucleotide polymorphisms or the loss and gain of novel gene families (Chaguza et al. 2019; Garud et al. 2019; Zhao et al. 2019 Groussin et al. 2021). However, to understand genome diversification across the range of bacterial strains with a species, it is necessary to model genome evolution over longer time scales. Recent work had a similar goal of understanding genome diversification in *Bacteroides* but relied on identifying core genes that were more similar to other bacterial species than to sequences of the bacterial species to which a strain belongs, thus biasing the detection of homologous recombination events. Another approach focused on detecting HGT events among bacterial species relied on identifying identical genomic regions, which limits detection to very recent HGT events between bacterial species (Groussin et al. 2021). Yet, it is unclear what fraction of the inter-strain differences in gene content these methods capture.

In this study, we trace the evolution of gene content among strains of multiple *Bacteroides* species through ancestral genome reconstruction in conjunction with a synteny-based approach to infer gene gain and loss events. By characterizing these events in multiple lineages of captive-ape-associated strains, we find that lineages of *B. xylanisolvens* have independently become enriched in genes related to metabolism of host mucosal glycans since their divergence with their closest human-associated strains.

## Results

The genome contents of the 131 *Bacteroides xylanisolvens* strains isolated from a range of host species and geographic locations vary substantially (Figure 1). The 39 *B. xylanisolvens* strains that we isolated from captive apes assort into three lineages—a Mixed-host lineage (within Clade A) containing 22 isolates collected from multiple species of great apes at the Houston and Columbus Zoos, and two other lineages represented by 4 isolates (Gorilla1 lineage within Clade A) and 13 isolates (Gorilla2 lineage within Clade B), all from gorillas at the Columbus Zoo (Figure 1). Over 90% of the *B. xylanisolvens* strains isolated from humans in industrialized countries fall into a single clade (Clade C) and are phylogenetically distinct from the strains isolated from captive apes, which are in Clades A and B. We also sequenced: (1) 12 *B. fragilis* strains isolated from orangutans at the Houston Zoo, which group together as a single clade and are closely related to a representative strain from humans (Supplemental Figure 1), and (2) three *B. ovatus* strains isolated from bonobos at the Columbus Zoo that form a monophyletic clade (Supplemental Figure 2).

**Figure 1.**
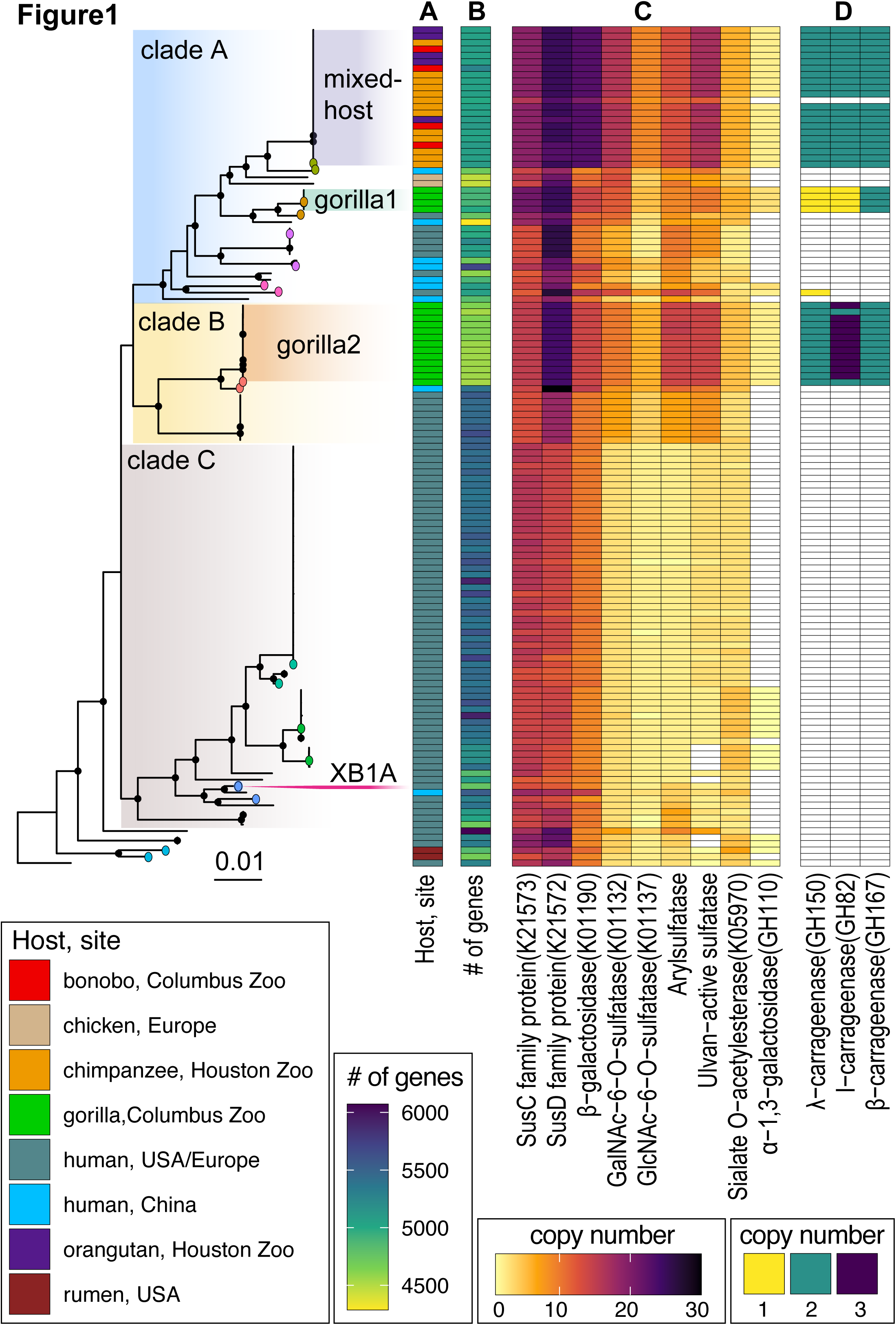
Core-gene phylogeny of *Bacteroides xylanisolvens*. Phylogeny of 131 strains generated with RAxML and based on 2098 genes present in >95% of genomes. Black dots denote nodes with >90% bootstrap support. Color-coded terminal nodes represent those clades used for phylogenetically independent contrasts with genomes being compared in the same color. Strains form three major clades (“Clade A”, “Clade B”, and “Clade C”), with strains isolated from captive apes assorting to three individual lineages within Clade A (“Mixed-host” and “Gorilla1”) and Clade B (“Gorilla2”). Phylogenetic position of species-representative strain XB1A (isolated from humans) is noted. To the right of the phylogeny, Column **A** indicates host species and sites from which each strain was isolated (with colors designated in left-most key) and the cells in Column **B** are shaded according to the number of predicted genes in the corresponding genome. Columns under headings **C** and **D** indicate copy numbers of those functional groups of genes that were acquired by captive-ape-associated strains since they diverged from their closest human-associated strains. (Functional classes under heading **C** are involved in mucin degradation, and functional classes under heading **D** are involved in carrageenan metabolism.)

### Convergent gene functional gain in three captive-ape lineages

By reconstructing gene gains and losses along the *B. xylanisolvens* strain phylogeny, a total of 64 functional groups show convergent increases in copy number in all three captive-ape lineages since they diverged from their closest human-associated strains (Supplemental Table 3). Genes with predicted functions related to mucin metabolism, including sulfatase and carbohydrate activity, exhibit some of the largest increases in gene copy numbers between captive-ape– and human-associated *B. xylanisolvens* strains (Figure 1). For instance, β-galactosidase (GH2, K01190) is involved in the degradation of the mucin backbone, and sialate O-acetylesterase (K05970) and α-1,3-galactosidase (GH110) remove side groups commonly attached to mucin (Larson et al. 1988; Corfield et al. 1992; Tailford et al. 2015; Raimondi et al. 2021). Both the SusC-like and SusD-like families of carbohydrate transporters, which are hallmarks of Polysaccharide Utilization Loci (PULs), are also convergently gained by all three captive-ape-associated lineages.

Because the functional group exhibiting the largest copy number differences (COG3119_NA_NA) contained several homologous gene groups (HGGs) that have different sulfatase predicted functions, we separated the HGGs into nine subgroups according to their Prokka annotations and looked for gene gains in each captive-ape-associated lineage (Mixed-host, Gorilla1 and Gorilla2). Both N-acetylgalactosamine-6-O-sulfatase (K01132) and N-acetylglucosamine-6-O-sulfatase (K01137) degrade the cap of mucin (enabling its catabolism), and both are convergently gained by representative strains in all three captive-ape-associated lineages. In addition, arylsulfatase and ulvan-activate sulfatase subgroups, which represent more general sulfatase functions, are also enriched in captive-ape strains. Finally, genes related to carrageenan metabolism are almost exclusive to strains isolated from captive apes: with the exception of a λ-carrageenase and a β-carrageenase gene observed in a single human isolate, λ-carrageenase (GH150), α-carrageenase (GH82) and β-carrageenase (GH167) genes are confined to *B. xylanisolvens* strains isolated from captive apes.

### Gene content evolution in *Bacteroides*

To determine whether the differences in gene contents observed in strains of *B. xylanisolvens* are typical of the *Bacteroides* genus as a whole, we examined variation in the number of predicted genes in 50 representative strains from *B. xylanisolvens*, *B. fragilis*, *B. thetaiotaomicron* and *B. ovatus*, most of which were isolated from human microbiomes (Figure 2A). We find that *B. fragilis* strains have significantly fewer genes relative to other *Bacteroides* species (Kruskal-Wallis all-pairs Dunn test, fdr correction, *p* < .001) (Figure 2A). In each species, the extent of gene contents differences increases with phylogenetic distance (Figure 2B; all *Bacteroides* species, Mantel test, *p* < .001); however, strains of *B. xylanisolvens* exhibit signficantly greater gene content divergence than either *B. fragilis* or *B. thetaiotaomicron* (but not *B. ovatus* (Figure 2B) (Supplemental Table 4) and relative to other *Bacteroides* species, *B. xylanisolvens* strains displays a greater differences in predicted gene numbers after correcting for phylogenetic relatedness (OLS, *p* = 5.6e-09).

**Figure 2.**
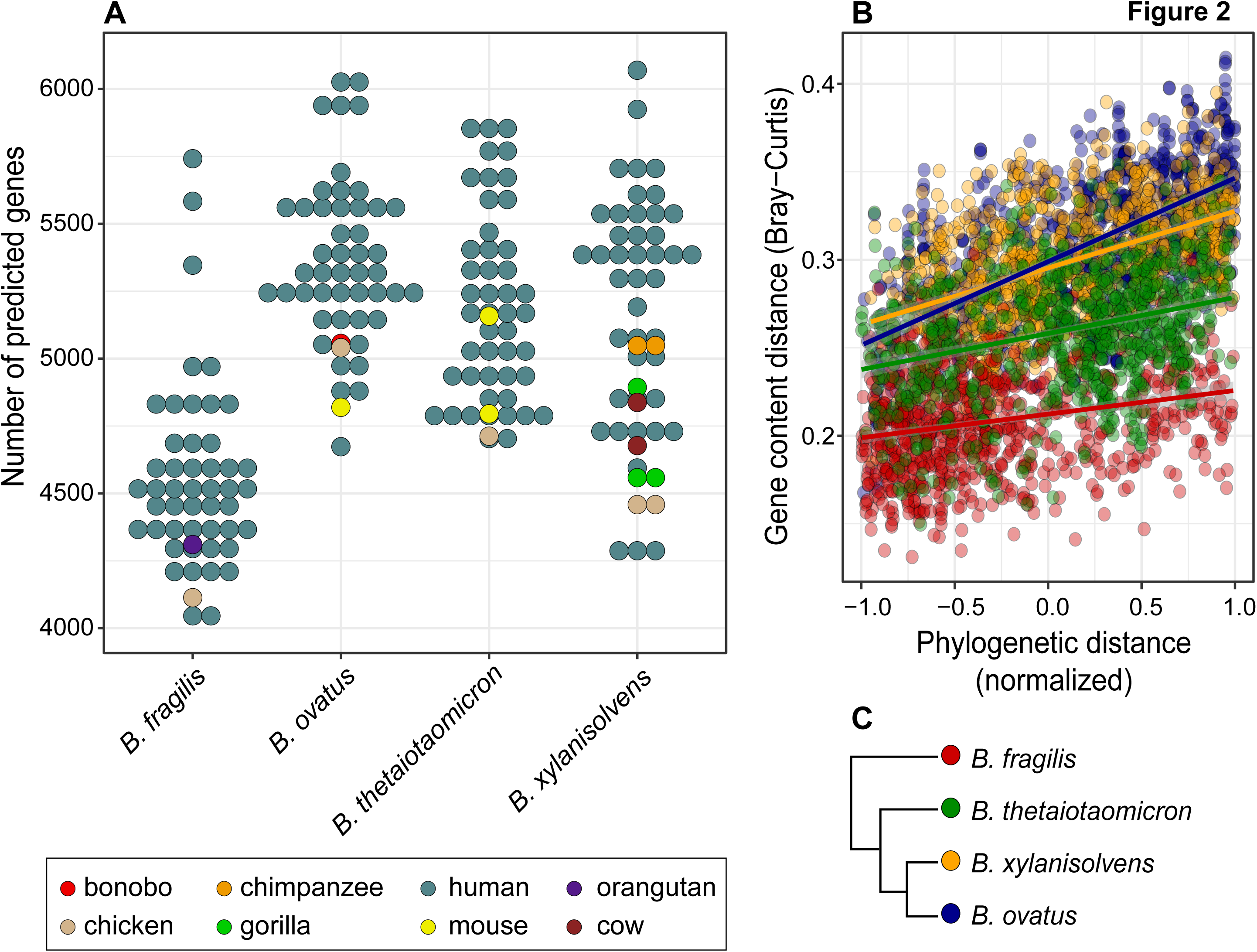
Genome content divergence across *Bacteroides* species. (A) Numbers of predicted genes in representative strains of *B. fragilis*, *B. thetaiotaomicron*, *B. ovatus*, and *B. xylanisolvens*. Data points are color-coded according to the host species from which strains were originally isolated. **(B)** Pangenome distance is positively associated with phylogenetic distance in representative strains of four *Bacteroides* species (Mantel test; *B. xylanisolvens*, *r* = .87, *p* = .0001; *B. ovatus r* = .80, *p* = .0001; *B. fragilis*, *r* = .33, *p* = .0007; *B. thetaiotaomicron r* = .60, *p* = .0001). **(C)** Phylogeny shows the phylogenetic relationships and color-coding of the four *Bacteroides* species.

### Gene gain and gene loss events in *B. xylanisolvens*

We investigated the relative frequencies of gene gain and loss events responsible for the divergence of gene contents in *B. xylanisolvens* across nine phylogenetically independent pairs of *B. xylanisolvens* strains (Figure 3). These comparisons included representatives from the three captive-ape-associated lineages and their closest human-associated strains, and six other contrasts of similar phylogenetic distance (Figure 1). By clustering genes that were lost or gained into events (*i.e*., segments of adjacent genes with a maximum gap of 5 genes to span ICE elements), we determined that events involving small segments (1–5 genes) represent the majority of both gain and loss events, and that the frequency of events decreases rapidly with size (Figure 3D). When gaps are prohibited, thereby limiting the analysis to strictly adjacent genes, the proportion of events over 50 genes in length is greatly reduced (Supplemental Figure 4). Only 15% of gene-gain events are attributable to duplication in that multiple copies are adjacent to one another. Infrequent large events account for a disproportionate amount of the genome variability (Figures 3E). Comparing the event sizes that represent 50% of the total number of genes gained (G50) or lost (L50), we find that gene-gain events are significantly larger (Kruskal-Wallis, *p* < .001) (Figure 3C). Other *Bacteroides* species also show gene gain events to be larger than gene loss events, with the majority of genes gained through events of at least 25 genes in length (Supplemental Figure 5), and captive-ape-associated strains tend to exhibit higher proportions of large gene gain events (Figure 3C).

**Figure 3.**
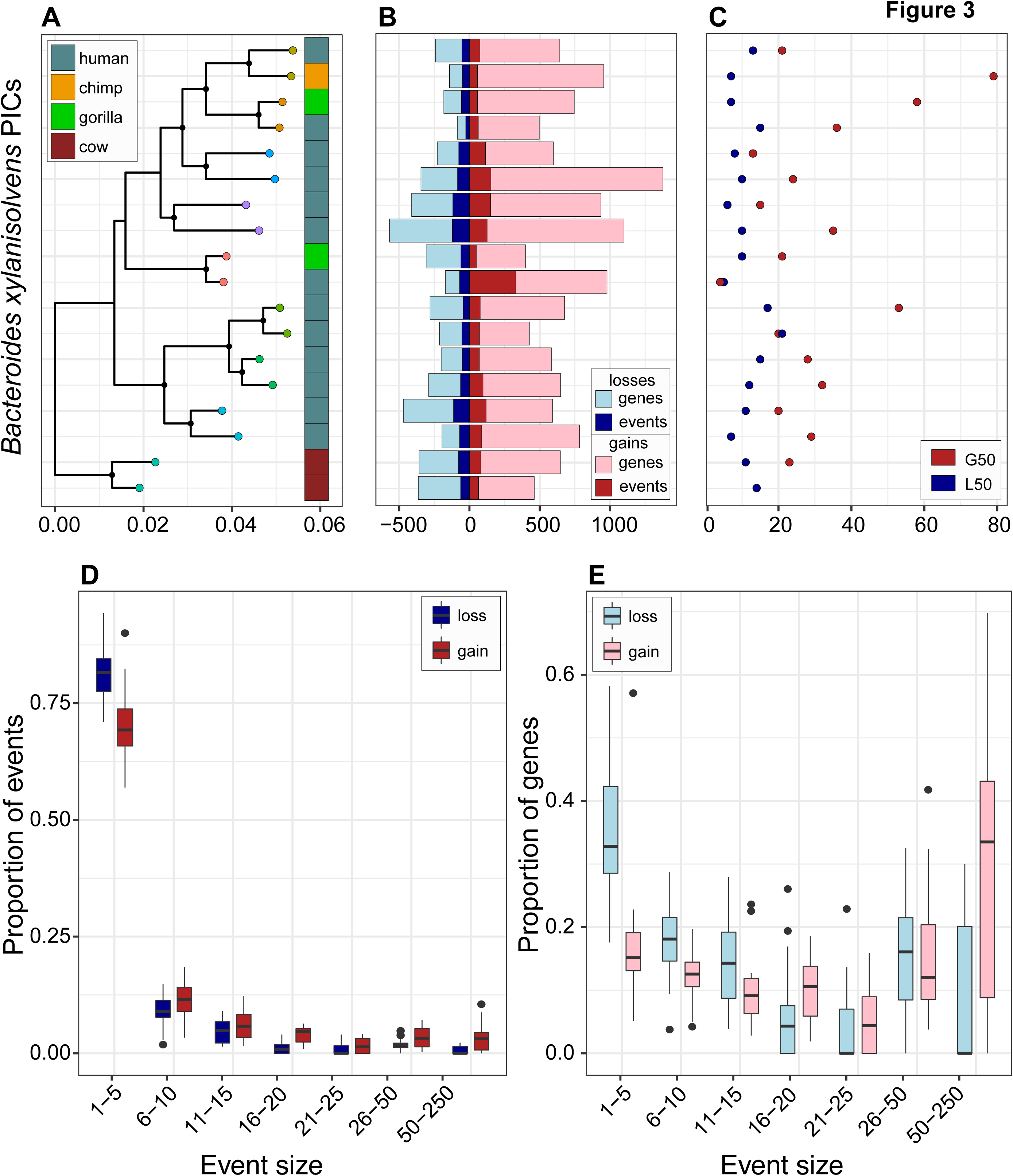
Gene gains and losses across *Bacteroides xylanisolvens*. (A) Phylogeny of the nine phylogenetically independent contrasts used to assess genome dynamics in *B. xylanisolvens* with pairs being compared in the same color. Column to the right of the tree indicates the host from which each strain was originally isolated. **(B)** Number of genes gained and lost, and the number of gain and loss events, accrued since a genome diverged from its sister taxa. **(C)** Event sizes at which 50% of genes are gained (G50) and lost (L50). **(D)** Distribution of the proportion of gain and loss events according to size, with boxplots displaying the median, IQR, and outliers (±1.5 IQR) across isolates within phylogenetically independent contrasts (*n* = 18). (E) Distribution of proportion of genes gained and lost according to event size with boxplots displaying the median, IQR, and outliers (±1.5 IQR) across isolates within phylogenetically independent contrasts (*n* = 18).

### Identifying potential source of genes gained by captive-ape-associated strains

To determine the relative contributions of within and between species transfer to these gene acquisition events, we identified highly similar copies (> 98% identity, > 98% alignment length) of genes gained by captive-ape-associated strains within the NCBI nr database. The largest fraction of gained genes represent those with highly similar copies in multiple *Bacteroides* species (Figure 4). Surprisingly, when considering those gained genes with highly similar copies restricted to a single species, both *B. thetaiotaomicron* and *B. ovatus* possessed larger numbers than *Bacteroides xylanisolvens*. A small fraction of genes gained by captive-ape-associated lineages exhibit taxonomic distributions beyond *Bacteroides*, with highly similar copies observed in *Parabacteroides, Prevotella* and other Bacteroidales genera, as well as other gut-associated taxa, including *Escherichia*.

**Figure 4.**
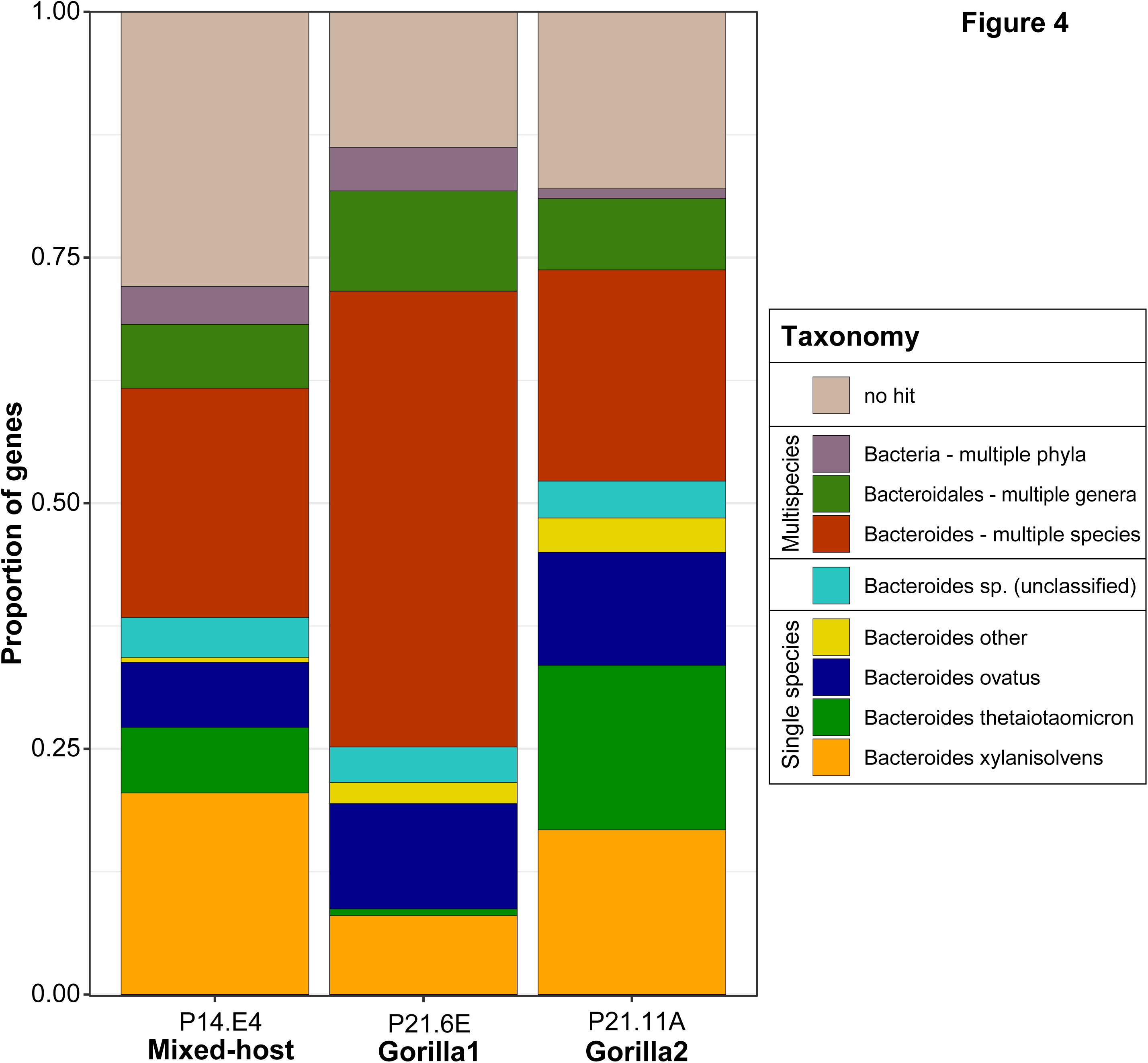
High-similarity copies of genes gained by captive-ape-associated lineages are found across multiple *Bacteroides* species. Proportion of genes gained by three representative isolates from captive-ape-associated lineages according to the taxonomic distribution of high similarity copies (>98% AAI) found in the NCBI nr database. Absolute numbers of genes gained by the three representative isolates are 956 (Mixed-host), 746 (Gorilla1) and 400 (Gorilla2).

### Independent acquisition of genomic segments by captive-ape strains

To determine whether polysaccharide utilization genes that are enriched in all captive-ape-associated lineages were acquired independently by each lineage, we focused on the 178 HGGs that were gained in at least two captive-ape-associated lineages and present in no more than three other hosts. Out of these 178 HGGs, 126 can be localized to a single region containing several Polysaccharide Utilization Loci (PULs) that are syntenic across multiple isolates (Figure 5). These PULs include SusC/SusD transporters, glycoside hydrolases with functions related to mucin and carrageenan metabolism, and sulfatase activity. Highly similar copies of many of the genes unique to captive-ape-associated lineages are observed in *B. ovatus* or *B. thetaiotaomicron* strains (Supplemental Table 5). Despite the overall conservation and synteny, strains from captive apes are variable in the presence of certain genes and segments within this region; *e.g*., isolates in the Gorilla1 lineage lack a span of genes containing sulfatase and glycoside hydrolase activity that are present in the other two clades.

**Figure 5.**
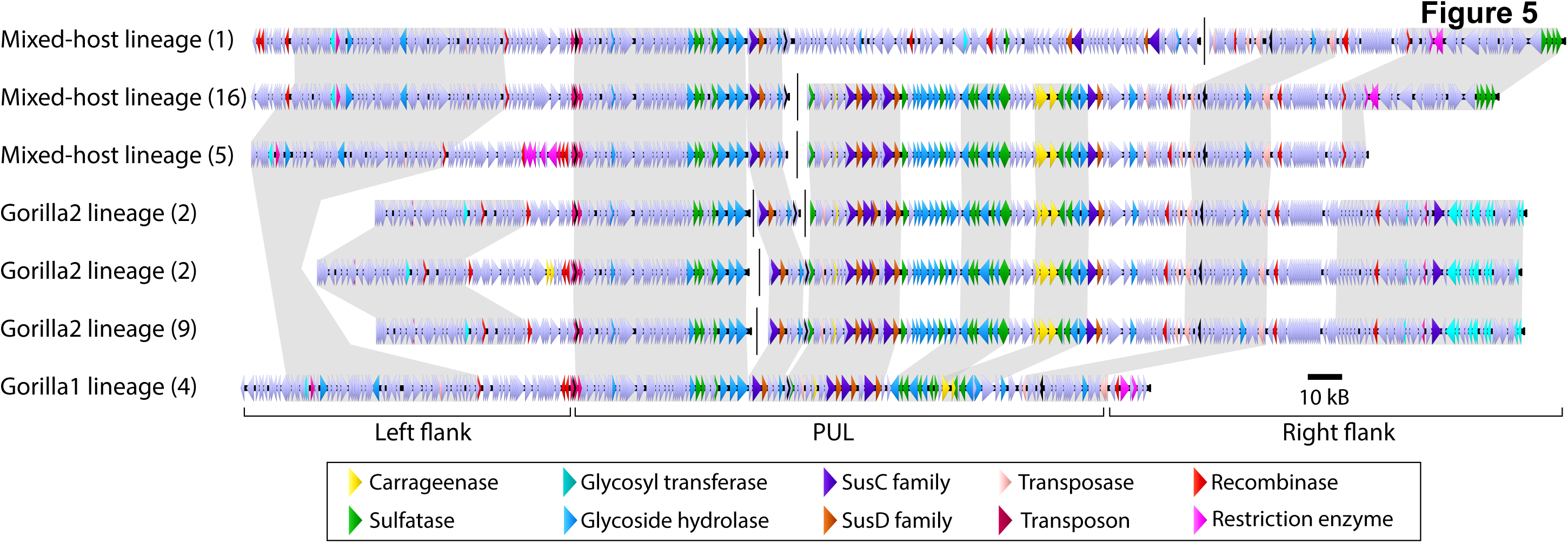
Organization of polysaccharide utilization loci (PUL) in *Bacteroides xylanisolvens* strains from captive apes. For each captive-ape lineage (“Mixed-host”, “Gorilla1”, “Gorilla2”; see Figure 1), alignments show PUL variants and flanking regions, with numbers of strains possessing a particular gene arrangement in parentheses. Genes are color-coded according to predicted function, as indicated in the key. To highlight similarities and differences among variants, grey sections link regions in which homologous genes are in the same order. Note that the left-flank regions in all Mixed-host variants are similarly divergent from those of all Gorilla2 lineages.

Several lines of evidence indicate that this region has been independently gained by multiple captive-ape-associated lineages: Firstly, HGGs belonging to this region are absence in other *Bacteroides xylanisolvens* isolates and loss by strains other than those from captive-ape-associated strains is unlikely. Next, the flanking regions of this PUL region differ between isolates belonging to the Mixed-host lineage in Clade A and the Gorilla2 lineage in Clade B, as evident in both the upstream and downstream sequences. (In isolates from the Gorilla1 lineage, the left-flanking region has partial synteny with the variants found in the Mixed-host lineage; however, the right-flanking region could not be mapped due to a contig break.) Finally, this span of genes is immediately flanked by mobile elements, which likely led to variable distributions among *B. xylanisolvens* strains.

### Sulfatase diversity and function

The PUL region in *Bacteroides xylanisolvens* can contains six sulfatase HGGs that are exclusive to strains isolated from captive apes and represents the largest proportion of gene functions gained by captive-ape-associated strains. On account of the enrichment in the overall number of sulfatase genes in captive-ape strains, we examined whether this trend also extends to the total diversity of sulfatase genes in captive-ape strains. A total of 91 HGGs present in at least 5 isolates were annotated as having arylsulfatase activity, of which 14 can be considered core genes due to their presence in captive apes and humans from all clades (Figure 6). However, eight sulfatase HGGs are found exclusively in the strains isolated from captive apes, four of which are present in all three lineages (Mixed-host, Gorilla1 and Gorilla2), and four shared by two of the three lineages (Mixed-host and Gorilla2). No sulfatase genes are exclusive to human-associated strains from different clades (*e.g.,* Clade A and Clade C).

**Figure 6.**
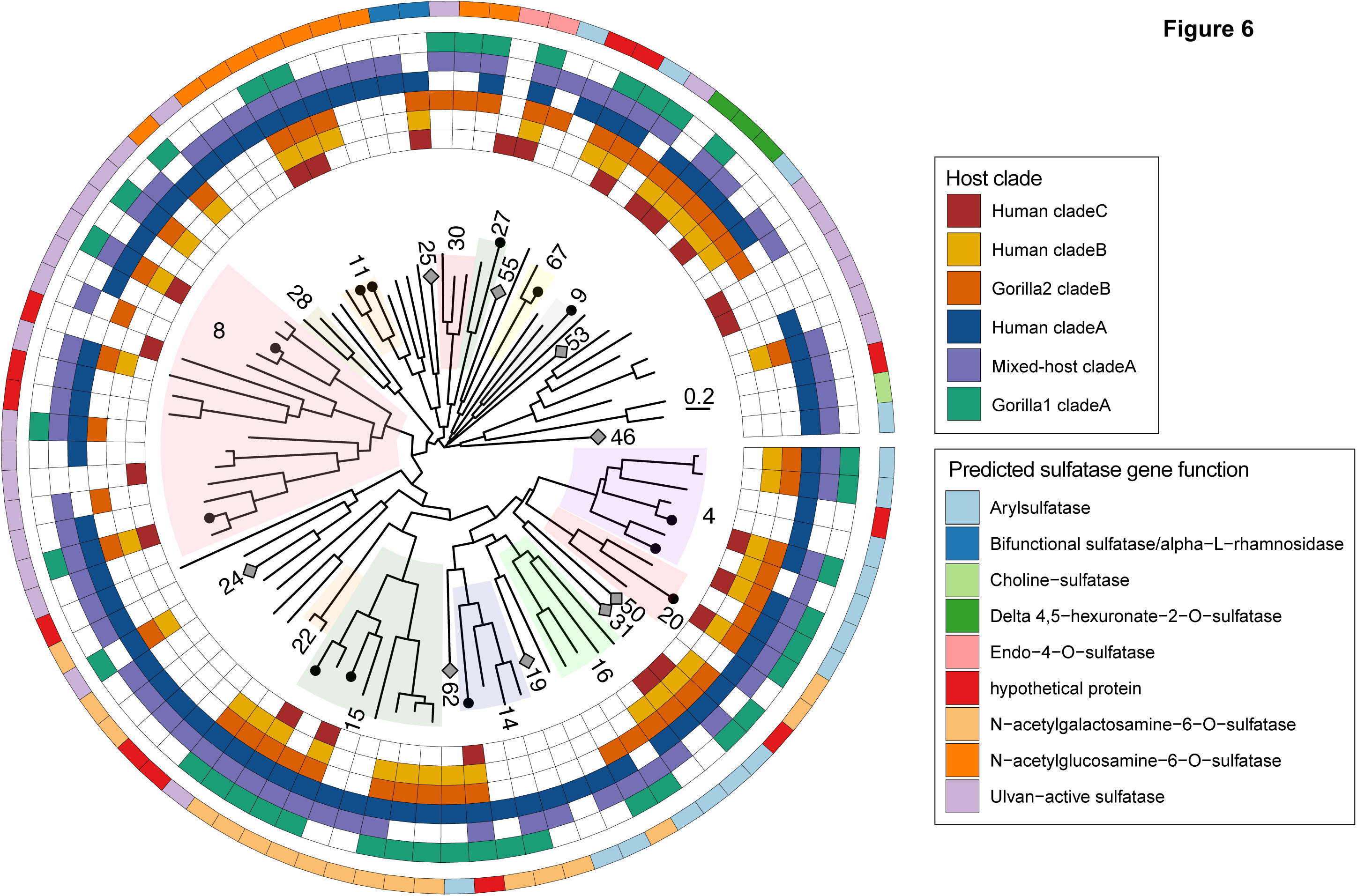
Phylogenetic relationships and distribution of sulfatase homologous gene groups. Clades are color-coded and/or numbered according to their S1 subfamily designations. Gray diamonds on terminal tips indicate subfamilies with only one representative, and black dots indicate the 14 core sulfatase genes detected in captive-ape and human-associated strains from all clades. The first six outer rings indicate the host distributions of each of the 91 sulfatase genes. The outermost ring is color-coded according to predicted gene function.

We observe large variation in the diversity of sulfatase genes in captive-ape-associated strains: Strains in Clade C, which includes the type strain XB1A as well as strains from humans residing in the US and Europe exhibit the lowest diversity, with only 23 sulfatase HGGs. In contrast, the human-associated strains within Clade A exhibit a total of 77 HGGs, and the Mixed-host lineage and Gorilla1 lineage in this clade possess 68 and 38 sulfatase HGGs respectively. In Clade B, there are 51 sulfatase HGGs in the Gorilla2 lineage, but only 38 in human-associated strains. Whereas strains from captive apes always have higher sulfatase gene copy numbers, they do not necessarily have higher gene diversity compared to strains from humans.

## Discussion

The gut microbiomes of great apes in captivity become populated with bacterial variants known only to occur in humans and are generally more similar in their overall microbiome composition to those in humans. To monitor the association between host gut environment differences and microbiome constituents, we examined the genomic changes that ensue in *Bacteroides* species whose representatives are present and prevalent in both humans and captive apes. Strains of *Bacteroides xylanisolvens* isolated from captive great apes in different locations independently acquired a large genomic region that confers gene functions related to host mucin and carrageenan metabolism. Whereas other studies have highlighted examples of gene transfer events between strains within or among *Bacteroides* species (Coyne et al 2014; Lange et al. 2016; Hussain et al. 2017; Singh et al. 2020), we assess both the extent and frequency with which transfers occur across *Bacteroides* species, showing that gene-gain events have played a disproportionate role in generating genomic diversity among strains: Roughly 50% of gene content differences between captive-ape-associated strains and their closest human-associated strains can be traced to events involving at least 25 genes, although such large events represent less than 10% of all events.

Since *Bacteroides* strains are known to gain and lose genes over short timescales (Groussin et al. 2021), it is possible that these PULs are transitory and represent neutral hitchhikers (Sousa et al. 2015). However, the independent acquisition of the homologous set of genes in multiple lineages, and the lack of any clearly inactivated genes in this region, suggests that they are functional. Several genes that are convergently enriched in captive-ape-associated strains and that are predicted to have a role in mucin metabolism have been shown to be upregulated when *Bacteroides xylanisolvens* D22 is grown on gastric mucin O-glycans (Pudlo et al. 2021). For example, one upregulated gene, that encoding sialate O-acetylesterase, removes mucin side groups, which enables cross-species foraging of mucin by pathogenic bacteria (Robinson et al. 2017). Additionally, ß-galactosidase, which shows a 150-fold increase in expression (Pudlo et al. 2021), is also known to be important in mucin degradation (Tsilford et al. 2015; Bell and Juge 2021).

Given the enrichment of mucin degradation genes, captive-ape-associated lineages of *Bacteroides* likely impact host-microbiome interactions. Whereas some studies have shown that bacteria that consume mucin can benefit host health (van Herreweghen et al. 2018; Kostopoulos et al. 2020), others suggest that increased degradation of mucin by *Bacteroides* can leave hosts vulnerable to colonization by opportunistic pathogens (Hickey et al. 2015; Fang et al. 2021). *B. xylanisolvens* strains may have acquired functions related to host mucin metabolism, due to the lack of fiber available in the captive-ape diet (Clayton et al. 2018). This situation is similar to what has been observed in *B. thetaiotaomicron*, which opportunistically metabolizes host-derived glycans when plant-derived glycans are absent from the diet (Bjursell et al. 2006). Moreover, the presence of additional carrageenan genes in captive-ape-associated strains may be a consequence of additives present in their diet, since carrageenan is ingredient in primate food sources (*e.g*., <u>Mazuri®</u>) recommended by the AZA (AZA Advisory Group 2017a,b,c) and an additive in processed foods (Weiner 2014). It is noteworthy that the ability for strains of various *Bacteroides* species, including *B. xylanisolvens,* to metabolize carrageenan (Li et al. 2017; Naimi et al. 2021) is attributable to HGT events originally from marine bacteria (Hehemann et al. 2012; Pudlo et al 2020).

The gut environment appears to promote horizontal gene transfer (Lerner et al. 2017); however, most methods used to infer HGT in the human gut—such as the presence of long stretches of nearly identical DNA in divergent organisms (Smilie et al, 2011; Brito et al 2016) or regional differences in base composition (Cortez et al. 2009)—overlook transfers among closely related species. Due to the high degree of sequence similarity in the genomes we appraised, we applied an approach that relied on parsimony and gene synteny to infer gene gain and loss events among strains. Whereas this method does not employ probabilistic models to quantify the likelihood of HGT of a given region, as applied by others (*e.g*., Adato et al. 2015; Sevillya et al. 2020; Cornet et al. 2021), it is a useful proxy to identify candidate regions of HGT among closely related strains. By this method, we determined the potential sources of HGT in *Bacteroides xylanisolvens* and find a large fraction of genes come from *B. ovatus* and *B. thetaiotaomicron*, perhaps not surprising given the close relationships and similar habitats of these species.

A limitation of our approach is that the ability to identify syntenic regions diminishes over time as subsequent insertion and deletion events erode synteny. Therefore, the sizes of gene gain and loss events can be underestimated due to successive interruptions. Conversely, there are also cases where a transfer event is followed by an internal duplication, which will both increase our appraisal of event sizes and underestimate the number of events required to generate the observed strain diversity.

The type strain of *B. xylanisolvens,* XB1A, harbors only a fraction of sulfatase genes observed in strains from other clades and is not representative of the diversity present in the most natural populations. For example, XB1A is not capable of growth on colonic mucin as a sole carbon source, because it lacks an enzyme belonging to the S1_20 subfamily that is responsible for explaining growth on colonic mucin (Luis et al. 2020), whereas many captive-ape-associated strains and human-associated strains harbor additional sulfatase genes belonging to this family S1_20 family. Acquisition of these new genes or loss of these genes by XB1A may indicate that some *B. xylanisolvens* strains can grow on colonic mucin; however, testing this is complicated by the fact that colonic mucin, unlike gastric mucin, is not readily available (Luis et al. 2020). Due to the wide variation in carbohydrate metabolism among closely related strains and the dependence on substrates related to particular host species and/or diet [78, 79], caution must be applied to narratives about the metabolic capabilities of a bacterial species based on few isolates (Pudlo et al. 2021; Luis et al. 2020; de Paepe et al, 2018).

Other studies have specified effective population size, habitat, lifestyle, selection and drift as factors affecting the evolution of bacterial pangenomes (Bobay and Ochman 2018; Maistrenko et al. 2020). Naturally, the frequency of HGT and the presence of molecular mechanisms that allow large gene gain events play a critical role (Wozniak and Waldor 2010; Brockhurst et al. 2019), and Groussin *et al*. (2021) report that *Bacteroides* species exhibit higher rates of new gene acquisition compared to *Bifidobacterium longum* and *Akkermansia muciniphila*. Additionally, Maistrenko *et al*. (2020) reported that two species of *Bacteroides* exhibit the highest levels of “genome fluidity”, computed as the ratio of unique gene families over the sum of gene families averaged over randomly chosen pairs of genomes from within a given species, of the 155 bacterial species analyzed. Therefore, it appears that *Bacteroides* exhibit some of the greatest differences in gene contents among all bacterial species surveyed, and our results show that even the colonization of new hosts prompted the gain of large genomic regions that expand metabolic properties beyond those present in their closest relatives.

## Materials and Methods

### Sample collection and bacterial strain isolation

Fecal samples of captive great apes from the Columbus and Houston Zoos (Supplemental Table 1) were collected using the BBL™ CultureSwab Collection and Transport System (BD Biosciences, NJ, USA), and shipped overnight at ambient temperature to our research laboratory where they were stored at 4°C. CultureSwabs were dunked repeatedly in 1 mL PBS to expel fecal material, and 100 µL of 10*^-3^*and 10*^-5^* dilutions were spread onto *Bacteroides* Bile Esculin Agar selective plates (Livingston et al. 2021). After 48 hours, approximately 16 colonies per sample were transferred into BHIS growth medium, passaged once (bacic and Smith 2008), and preserved in 25% glycerol. DNA extractions were performed using phenol-chloroform-isoamyl alcohol (25:24:1) (Goodman et al. 2011), and strain identities determined by amplifying and sequencing a portion of the Gyrase B gene. Strains with different Gyrase B sequences were selected for whole genome sequencing using the SWIFT 2S™TURBO kit (Swift Biosciences, MI, USA) on the Illumina NovaSeq PE151 platform, yielding a total of 39 *Bacteroides xylanisolvens* strains, 3 *B. ovatus* strains and 12 *B. fragilis* strains.

### Genome assembly

Raw fastq read qualities were visualized using FastQC (Andrews 2010) with read-trimming and quality-filtering implemented in the PE mode of trimmomatic (Bolger et al. 2012). *De novo* assemblies were generated with SPAdes in isolate mode with error-correction (Bankevich et al. 2012). Taxonomy of contigs was determined with centrifuge (Kim et al. 2016), which recognized very low levels (<0.01% of reads) of human DNA contamination. Only contigs >1,000 bp in length with >10x coverage and having a centrifuge taxonomic assignment to *Bacteroides* were retained. Sspacer and gapfiller were used to close gaps (Boetzer et al. 2011; Nadalin et al. 2012). Contigs were aligned and genome coverage assessed with bbmap from bbtools (Bushnell 2014), completeness and contamination evaluated by CheckM (Parks et al. 2015), and taxonomic assignments determined with GTDBTk (Chaumeil et al 2020) (Supplemental Table 1).

### Filtering publicly available genomes

Genomes typed to *Bacteroides*, including *B. xylanisolvens* (*n* = 92), *B. ovatus* (*n* = 92), *B. fragilis* (*n* = 170), and *B. thetaiotaomicron* (*n* = 74) were downloaded from the NCBI Prokaryotic Genomes server. GTDBTk and CheckM were run on each genome to validate strain taxonomy, and to ascertain completeness and contamination. Applying thresholds of >95% completeness and <5% contamination removed 23 downloaded genomes from subsequent analyses (Supplemental Table 1 & 2). Completeness and contamination thresholds were determined empirically using *B. xylanisolvens* assemblies and were selected to ensure that the number of predicted genes was not significantly associated with genome contamination (*t*(122) = 1.00, *p* = .32) or completeness (*t*(122) = –1.94, *p* = .055).

### Genome annotation and pangenome construction

*Bacteroides* genomes were annotated with Prokka (Seeman 2014). We ran Roary (Page et al. 2015) to determine gene presence across strains belonging to each *Bacteroides* species. For an individual species, three genomes of a closely related species were used as an outgroup; for instance, three genomes from *B. ovatus* were used as the outgroup for the Roary run of *B. xylanisolvens*. Because Roary typically assigns orthology based on synteny, we used the no-split-paralogs flag to generate homologous groups of genes (HGG) rather than strictly orthologous groups of genes to identify convergently acquired genes having different neighboring genes. Representatives from each HGG were annotated using a suite of databases including SulfAtlas (Barbeyron et al. 2016), eggNOG database (Huerta-Cepas et al. 2019), KEGG KofamKOALA (Aramaki et al. 2020), and CAZy database (Cantarel et al. 2009). For each *Bacteroides* species, a phylogeny based on genes present in at least 95% of stains was generated with RAxML, implementing the GTRGAMMA model and 100 bootstraps (Stamatakis et al. 2014).

### Reconstructing ancestral genome content and predicted gene functions

Gene gains and losses were reconstructed along the strain phylogeny of eacg *Bacteroides* species by applying Wagner parsimony with a gain/loss penalty of 2 (Pál et al. 2005; Csurös 2010; Zaremba-Niedzwiedzka et al. 2013; Oyserman et al. 2016), which, unlike other values tested, yielded no significant difference in the number of genes between ancestral and extant genomes (Kruskal-Wallis test, *p* = .289).

Homologous groups of genes (HGGs), as identified by Roary, were clustered into larger functional groups based on their eggNOG classification, KEGG K number, and Carbohydrate-Active EnZyme modules (*e.g*., COG3250@2 | Bacteria_K01190_GH2 *vs*. COG5434@2 | Bacteria_NA_GH82, where NA indicates no annotation was made). Due to the independent origins of bacterial lineages within captive-ape, we consider functional groups to be “convergently enriched” if there are increases in copy numbers in all three captive-ape-associated lineages (Mixed-host, Gorilla1 and Gorilla2) since they diverged from their respective common ancestors with their closest human-associated strains. Even though we include only those strains that are at least 95% complete, some instances of gene gain or loss might be missed in subsequent analyses; however, if gaps occur at random, as suggested by the distribution of shared genes, the occurrence and inference of parallel changes in different lineages are unlikely to occur by chance.

### Gene content distance

To examine gene-content differences across a broad phylogenetic scale, we selected 50 representative strains of each species based on the hierarchical clustering of phylogenetic distances applying a *k* = 50 parameter. The degree of dissimilarity between gene repertoires of strains was calculated from the Bray-Curtis distances of the gene presence-absence matrix produced by Roary, in which columns are isolates, rows are HGGs, and values are gene-copy numbers. Because pangenomes and phylogenies were generated individually for each *Bacteroides* species, phylogenetic distances were normalized by subtracting the mean phylogenetic distance for all strains in a given species and dividing by the standard deviation. To evaluate whether differences in gene contents are associated with phylogenetic distance, we applied Mantel tests between phylogenetic distance and gene content distance, with fdr-corrected *p*-values.

### Gene content evolution among *Bacteroides* species

Because pairwise comparisons between strains are not phylogenetically independent, we evaluated gene content evolution among phylogenetically independent contrasts (PICs) of strains. We used an in-house script that maximized the number of PICs at a specified phylogenetic distance between strains. For *B. xylanisolvens,* we applied a cutoff value of .008 because this yielded three PICs between captive-ape-associated strains and their most closely-related human-associated strains and six additional PICs between human-associated strains at comparable phylogenetic distance. Because this cutoff is approximately 2.2 standard deviations below the mean phylogenetic distance among all pairs of the 50 representative *B. xylanisolvens* strains, we applied a cutoff of 2.2 standard deviations below the mean phylogenetic distance for other *Bacteroides* species. This generated 7–17 PICs in these other *Bacteroides* species (Supplemental Figures 1-3).

For each PIC, we used output from Count to identify the genes that are either gained by strain 1, lost by strain 1, gained by strain 2, or lost by strain 2 since their MRCA. We excluded the small fraction of HGGs where the gene copy number of the MRCA was not the same as the copy number in either strain 1 or strain 2. For each gene category, we iterated through GeneIDs based on their position in the gff file, and when the subsequent GeneID was contained within the same contig and within five genes of the current GeneID, we considered them as originating from the same genomic event. (The threshold window of five genes enables segments of adjacent genes to span an interruption of a few genes that were frequently annotated as integrative and conjugative elements.) Analyses were also conducted for cases in which adjacent genes were considered part of the same gain or loss event. Although most of the copy-number variation of HGGs is due to their presence in one genome and absence from another, occasionally strains possess the same HGG in different copy numbers. In such cases, it is often not possible to determine which of the additional copies were gained or lost, so we selected the gene copy residing on the largest genomic event in order to minimize the number of events required to explain the gene content differences. To determine the probable source of gained HGGs, we performed a DIAMOND blastp search of the NCBI nr database and filtered out hits having >98% identity and >98% alignment length. These cutoffs were chosen to identify hits to other bacterial species were comparably similar to the average percent identity and alignment length of hits to *Bacteroides xylanisolvens* strains. Gene gain events can result from HGT from an external source, by intragenomic duplication events, or from a combination of the two. We considered gene gain events to result from duplication events, as opposed to HGT, when all HGGs in an event are represented by two or more copies.

### Identifying regions of independently acquired genes

To search for genomic regions that were independently acquired by multiple captive-ape-associated lineages, we identified HGGs that were gained by at least two of the three captive-ape-associated lineages (Mixed-host, Gorilla1, and Gorilla2) since their MCRA with their closest human-associated strain and present in fewer than three strains from other hosts. We extracted regions containing these HGGs, along with 100 upstream and downstream genes, and used Geneious to align, annotate, and visualize regions among strains.

### Distributions and functions of sulfatase genes

Protein sequences representing sulfatase HGGs (*e.g*., those annotated as COG3119) that were present in at least five *B. xylanisolvens* isolates were aligned in mafft (Katoh and Standley 2013) and subjected to phylogenetic reconstruction using fasttree (Price et al. 2010). Sulfatase subfamily designations were based on Sulfatlas annotation, and gene functions were predicted from Prokka annotations.

## Supporting information

Supplementary Figures S1 through S5

Supplementary Tables S1 through S5

## Acknowledgements

The authors would like to thank the following people and organizations for providing fecal samples from captive primates: Judy McAuliffe and Houston Zoo staff, Audra Meinelt, Laura Pierson and Columbus Zoo staff, and Project Chimps staff. We also thank Jen Lavoie and Angela O’Donnell for their assistance in sequence library preparation, and for discussions and comments on the manuscript.

## Data and code availability

Genome assemblies of strains sequenced in this study are available on NCBI under the BioProject number PRJNA767094. Code used in data processing and analysis available through the github repository https://github.com/ahnishida/Bxy_strains.

