## Supplementary Figures S1 through S5 for "Origins and Evolution of Novel *Bacteroides* in Captive Apes"

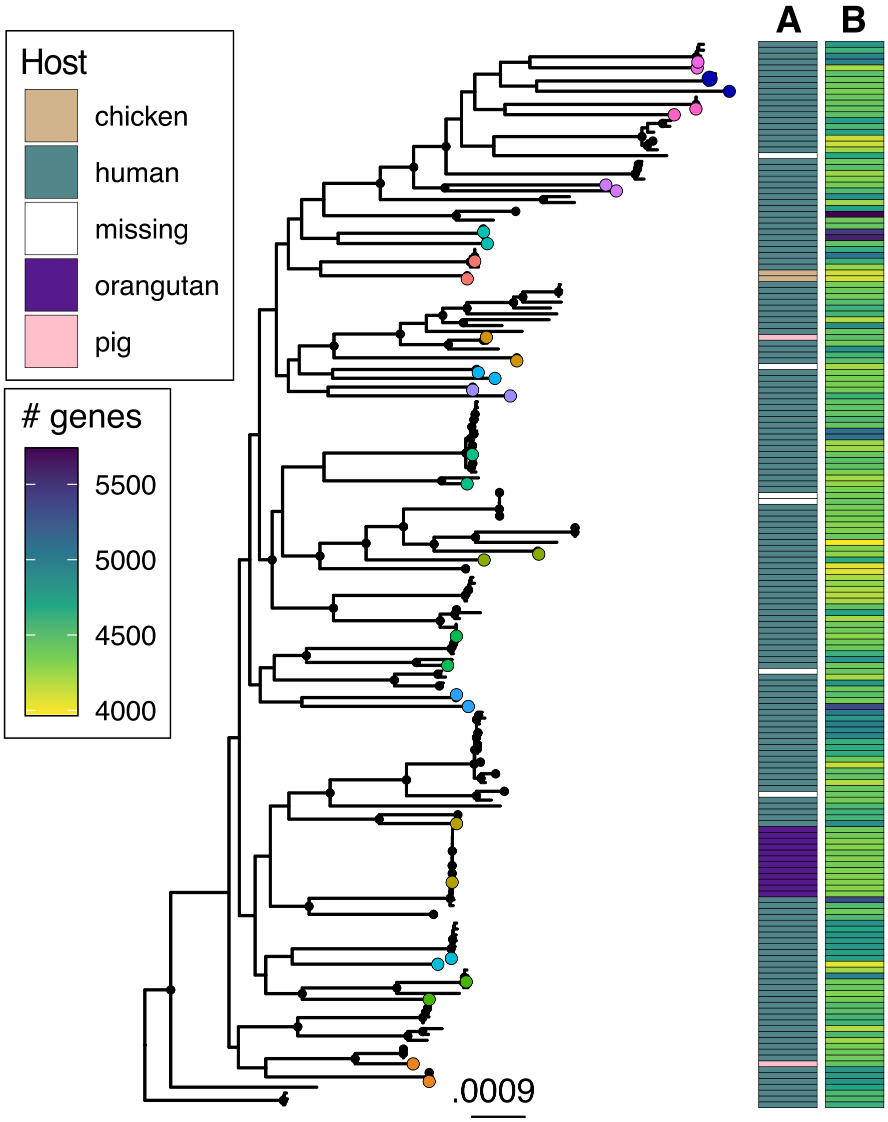


**Supplemental Figure 1. Core-gene phylogeny of 182 *Bacteroides fragilis* genomes.** Black dots denote nodes with >90% bootstrap support. Color-coded terminal nodes represent those clades used in phylogenetically independent contrasts, with genomes being compared highlighted in the same color. Cells in Column A are shaded to indicate host species from which strains were isolated, and cells in Column B are shaded to indicate the number of predicted genes in the corresponding genome.

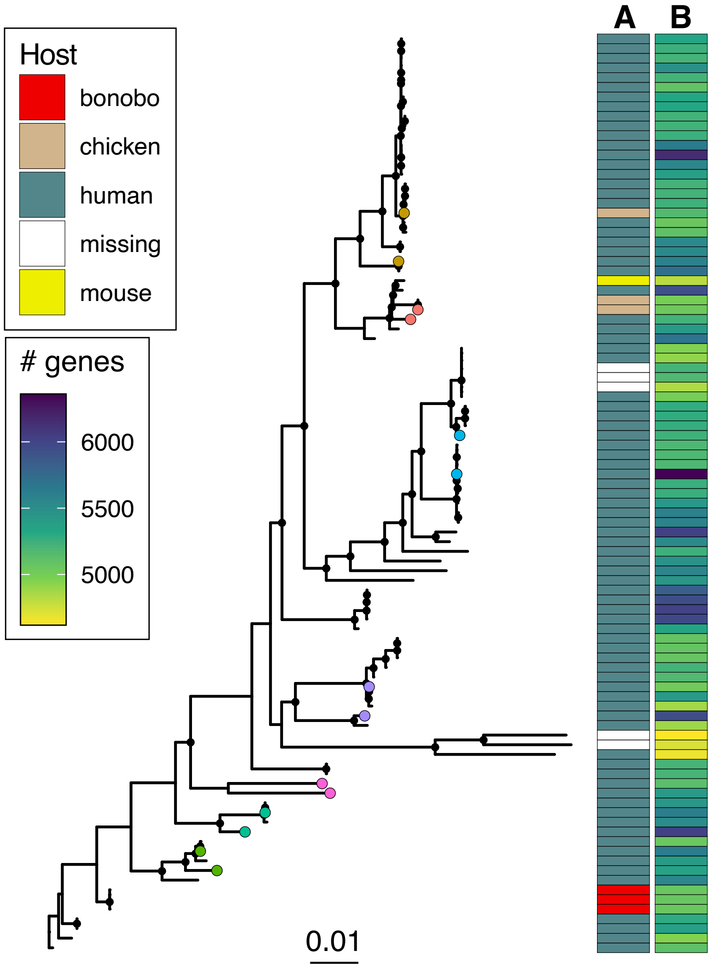


**Supplemental Figure 2. Core-gene phylogeny of 95 *Bacteroides ovatus* genomes.** Black dots denote nodes with >90% bootstrap support. Color-coded terminal nodes represent those clades used in phylogenetically independent contrasts, with genomes being compared highlighted in the same color. Cells in Column A are shaded to indicate host species from which strains were isolated, and cells in Column B are shaded to indicate the number of predicted genes in the corresponding genome.

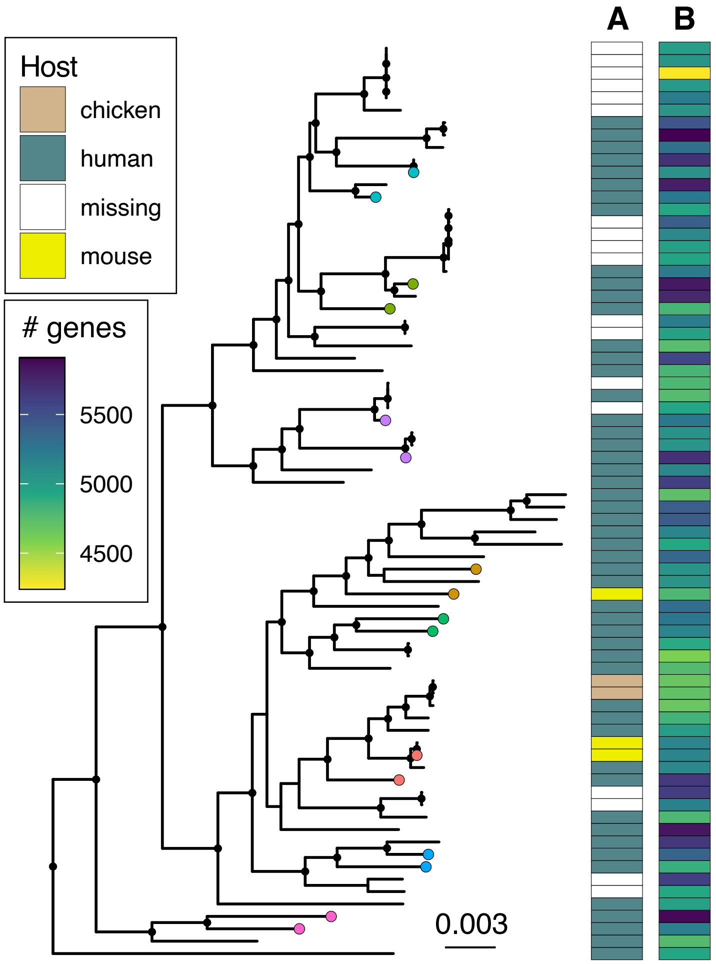


**Supplemental Figure 3. Core-gene phylogeny of 74 *Bacteroides thetaiotaomicron* genomes.** Black dots denote nodes with >90% bootstrap support. Color-coded terminal nodes represent those clades used in phylogenetically independent contrasts, with genomes being compared highlighted in the same color. Cells in Column A are shaded to indicate host species from which strains were isolated, and cells in Column B are shaded to indicate the number of predicted genes in the corresponding genome.

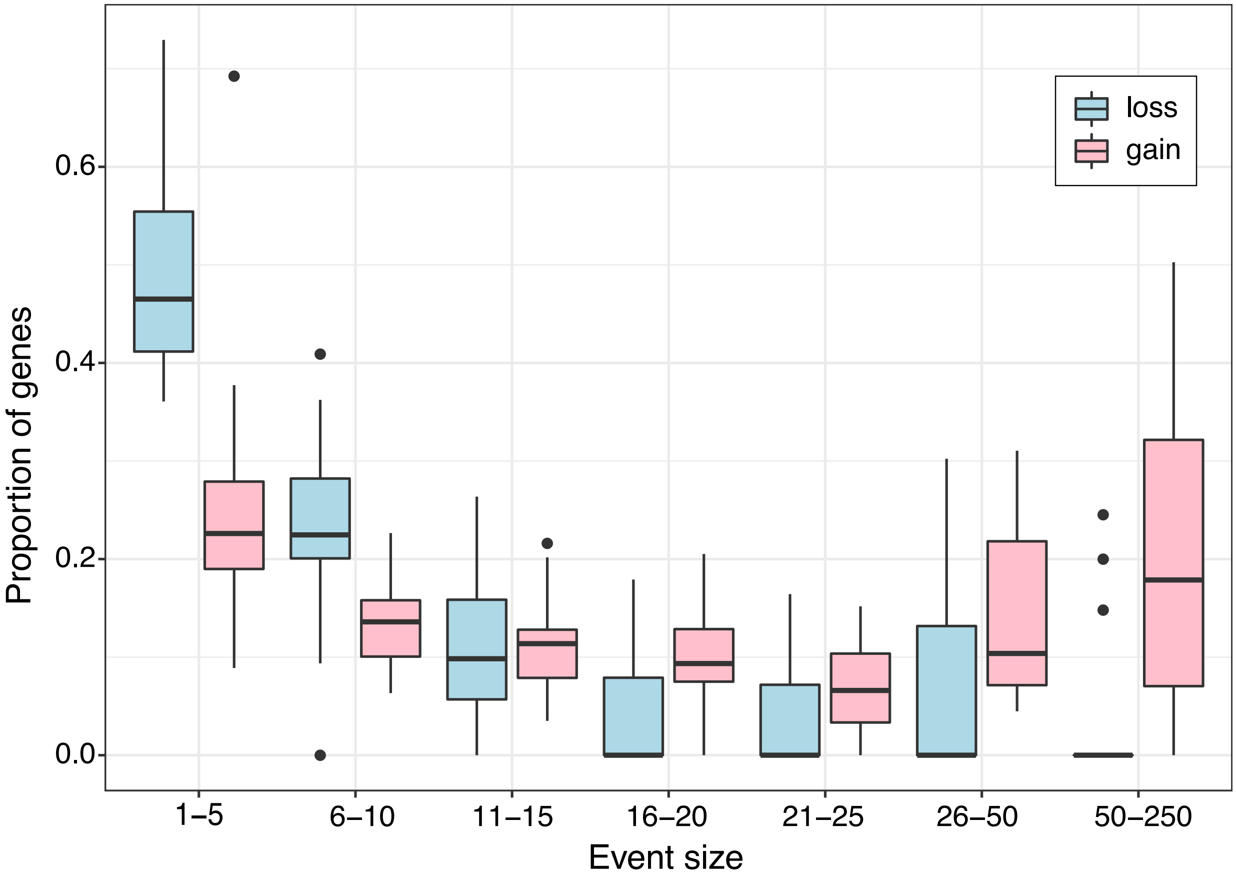


**Supplemental Figure 4. Proportions of genes gained and lost in relation to event size.** Boxplots display median, IQR, and outliers (±1.5 IQR) across isolates used in phylogenetical independent contrasts (*n* = 18) when events are limited to strictly adjacent genes.

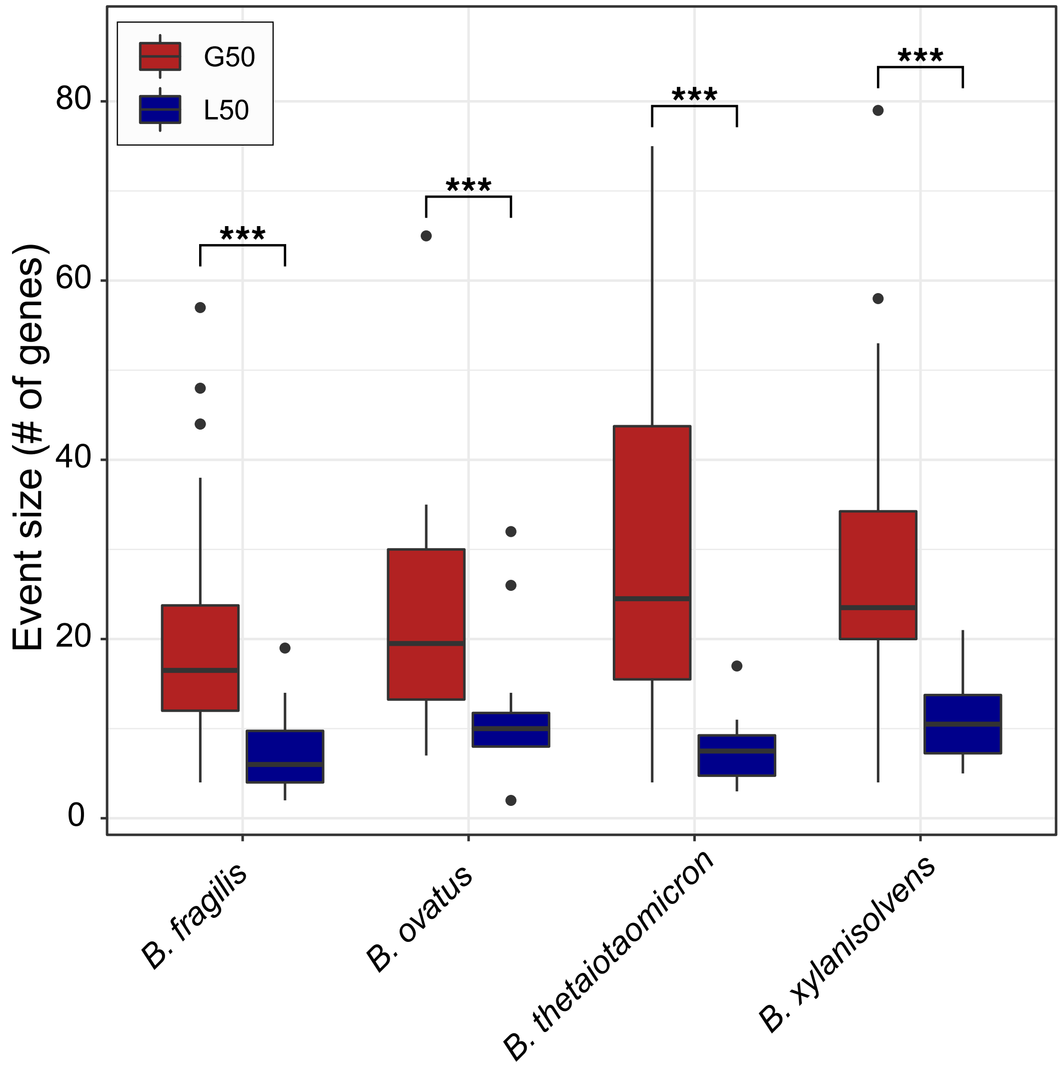


**Supplemental Figure 5. Gene gain and loss across Bacteroides species.** Shown are sizes of events at which 50% of genes are gained (G50) or lost (L50) computed for isolates used in phylogenetic independent contrasts. G50 is significantly larger than L50 in each of the *Bacteroides* species considered (Kruskal-Wallis, all comparisons, $p < .001$).
